# TSdb: A Curated Database for Terpene Synthases and Their Application in Natural Product Mining

**DOI:** 10.1101/2025.09.01.673588

**Authors:** Tongyu Gu, Dengming Ming

**Affiliations:** Nanjing Tech University

**Keywords:** terpene synthase, database, biosynthetic gene clusters (BGCs), protein function annotation

## Abstract

Terpenoids constitute nature’s most chemically diverse metabolite family with vital pharmaceutical and industrial applications, yet existing databases lack systematic integration of precursor metabolic enzymes (HMGR, DXS) and mechanistic insights into terpene diversification. To bridge this gap, we developed the Terpene Synthase Database (TSDB), distinguishing itself through three key innovations: (1) comprehensive integration of MVA/MEP pathway enzymes with downstream terpenoid synthases, (2) enhanced functional annotation via InterProScan domain mapping and phylogenetics to decode catalytic plasticity, and (3) unprecedented taxonomic breadth spanning 456,142 non-redundant sequences across 30,491 taxa. By consolidating data from BRENDA, UniProt, TeroKit, and ocean gene clusters through rigorous BLASTp deduplication (95% identity cutoff), TSDB reveals 3,499 Gene Ontology terms highlighting core functions like isoprenoid biosynthesis (GO:0019288) and metalloenzyme catalysis. Validation against MIBiG gene clusters (e.g., BGC0001324) demonstrates precise identification of terpene cyclases, P450 monooxygenases, and prenyltransferases with residue-level active site annotations. As the first resource connecting precursor metabolism to structural diversity, TSDB enables accurate gene-enzyme-product prediction for enzyme engineering and natural product discovery.

## 1 Introduction

Terpenoids represent the most structurally diverse and functionally versatile family of natural products. Their broad biological activities and ecological roles make them central resources for drug development, agricultural protection, and industrial applications. In medicine, sesquiterpenoids such as artemisinin have revolutionized malaria treatment strategies [1], while diterpenoids like paclitaxel serve as critical sources of anticancer agents [2]. Monoterpenes and sesquiterpenes mediate ecological interactions through plant volatiles [3] and demonstrate potential in biofuel development [4].

The structural diversity of terpenoids stems from the complexity of their biosynthetic pathways. In eukaryotes (e.g., plants and fungi), the mevalonate (MVA) pathway initiates with acetyl-CoA to generate isopentenyl pyrophosphate (IPP) and its isomer dimethylallyl pyrophosphate (DMAPP), which are precursors for sterols and certain terpenoids. In contrast, the methylerythritol phosphate (MEP) pathway in bacteria, plant plastids, and some protists utilizes pyruvate and glyceraldehyde 3-phosphate to produce IPP/DMAPP, primarily driving the synthesis of monoterpenes, diterpenes, and carotenoids.

Terpenoid biosynthesis begins with the condensation of IPP and DMAPP, followed by cyclization catalyzed by terpene synthases (TSs). Subsequent modifications by cytochrome P450 oxidases (P450s), glycosyltransferases (UGTs), and other enzymes introduce functional groups such as hydroxyl and glycosyl moieties [5]. Notably, TSs and P450s exhibit remarkable catalytic plasticity during evolution [6], where a single enzyme may catalyze multiple reactions, while structurally similar enzymes may yield distinct products. This substrate promiscuity and functional divergence expand the chemical space of terpenoids but complicate the precise prediction of gene-to-product relationships, especially in the absence of comprehensive enzyme annotations and structural data.

Advances in genomics and metabolomics have accelerated the discovery of TS genes. However, existing databases such as UniProt, KEGG, and BRENDA provide fragmented information with limitations in data integration, annotation depth, and update frequency. Specialized TS databases remain scarce. The TeroKit database (http://terokit.qmclab.com/index.html), developed by Prof. Ruibo Wu’s team in 2020, integrates 180,000 terpenoid compounds (TeroMOL)[7] and 13,000 enzymes (TeroENZ)[8], focusing on terpene backbone assembly and late-stage modifying enzymes (e.g., P450s). However, it lacks systematic coverage of precursor metabolic enzymes (e.g., HMGR and DXS in MVA/MEP pathways) and mechanistic insights into catalytic specificity.

We constructed the Terpene Synthase Database (TSDB) to address these gaps by integrating multi-source data from BRENDA, UniProt, TeroKit, and marine microbial gene clusters. TSDB comprises 456,142 non-redundant enzyme sequences spanning 30,491 taxonomic units, linking precursor metabolism with terpenoid diversification. Enhanced functional annotations via sequence analysis, InterProScan domain prediction, and phylogenetic analysis establish TSDB as a high-quality platform for terpenoid biosynthesis research. This resource supports elucidating TS roles across pathways and provides foundational data for enzyme engineering, metabolic regulation, and natural product discovery.

## 2 Methods

### 2.1 Construction of a comprehensive terpene synthase database

#### 2.1.1 Acquisition of terpene synthase data

##### 2.1.1.1 Terpene synthase data derived from Brenda

Tiangang Liu[9] et al. specifically described the enzymes required for terpenoids in MVA and MEP biosynthetic metabolic pathways in “The Construction of Terpenoids Efficient Synthesis Platform and Terpenoids Product Batch Mining.”

We have previously screened enzyme sequences using the text search functions of BRENDA and UniProt, so that substances involved in the synthesis of terpenoid precursors in the MEP and MVA pathways were obtained. We extracted essential details from this process, including the BRENDA ID, UniProt ID, names of terpene synthases, EC number, organism source, and information from Swiss-Prot/TrEMBL. This information has been compiled and deposited into a FASTA file. Based on that, this study utilized BRENDA’s API interface and Python scripts to grab the newly added enzyme sequence data in recent years from the BRENDA database based on the EC numbers in this FASTA file, thus further refining and updating the data content and the sequence richness of the comprehensive database of terpene synthases. The final total was 14 classes of synthases, 17 EC numbers, and 364,066 sequences. We call these Collection A.

##### 2.1.1.2 Processed data originating from Uniprot and Terokit

UniProt (https://www.uniprot.org/) is a comprehensive database of protein sequence and function information widely used in life science research. It provides essential support in protein function and study. UniProt contains three main components: UniProtKB (containing annotatable protein sequences), UniRef (clustering data for fast similarity searches), and UniParc (providing non-redundant protein sequence records). With UniProt, users can access protein function, structure, interactions, modifications, and related biology information.

TeroKit (http://terokit.qmclab.com/) [10] is a database focused on terpenoid compounds, covering information on the structure, function, and biosynthetic pathways of terpenoid-derived compounds, including terpene synthases (TSs), steroids, and their derivatives. It also organizes enzymes involved in terpenoid biosynthesis and provides links to external databases for enzyme species origin, sequence, family, and protein structure. Although TeroKit offers a wealth of enzymatic data, it covers limited information on the functional aspects of terpenoid compounds, primarily focusing on enzymes involved in synthesizing terpene scaffolds and chemical modification reactions.

This study builds upon the previously integrated data from our research group, further expanding and updating the terpene synthase information obtained from these two databases. The research group had previously processed and integrated terpene synthase data sourced from UniProt and TeroKit, ultimately aligning 62,378 sequences, which were then saved as FASTA format files. To facilitate subsequent annotation calculations, all terpene synthase data were divided into two categories: 9,937 sequences that were manually annotated and labeled as “reviewed” by UniProt, and 9,937 sequences that were not manually annotated and labeled as “unreviewed,” totaling 52,282 sequences.

It should be noted that although 62,378 sequences are included, about a hundred IDs failed to be matched in the UniPort. Hence, the number of entries in the file is not exactly equal to the sum of the sequences in the two categories mentioned above.

##### 2.1.1.3 Data derived from marine microbial gene clusters

We also extracted relevant terpenoid data from the Ocean Microbiomics Database (https://microbiomics.io/ocean/). The OMD database provides detailed information on marine microbial gene clusters, and users can export the appropriate data in Excel format. The exported information includes BGC, Genome, Taxonomy, BGC length, BGC complete, BGC representative, Genome Clusters of Orthologous Groups (GCF), Genome Cluster Count (GCC), products, Non-Ribosomal Peptide Synthetases, Type I Polyketide Synthases, Type II/III Polyketide Synthases, RiPPs, and terpenoid compounds, among others. [11]

To further obtain sequence data on terpene synthases, we downloaded GBK format files of biosynthetic gene clusters (BGCs) containing detailed sequence information from the database. A total of 51,861 gene cluster entries were included, focusing on the CDS (coding sequence) regions, which also contained terpene synthase sequences related to terpene compound biosynthesis. The terpene synthase sequence files were obtained by screening marine microbial gene clusters using the keyword “terpene” and by writing a Python script to extract terpene synthase sequences from the CDS (coding sequence) regions. These sequences were then organized into a file named bgc_CDS_ts and saved in FASTA format. Subsequently, makeblastdb and blastp commands were used to align these sequences with terpene synthase data integrated from UniProt and TeroKit, identifying 31,259 sequences.

##### 2.1.1.4 Data Screening and Integration

This study performed data deduplication and sequence alignment on the collected sequence information to establish a high-quality and accurate database query system.

NCBI’s BLAST (Basic Local Alignment Search Tool) was employed as the core data deduplication and sequence alignment method. BLAST is a fundamental bioinformatics tool widely used for gene annotation, functional prediction, species identification, and other tasks [12]. Similar sequences can be identified by aligning nucleotide or protein sequences with known databases and potential biological relationships revealed.

Based on the type of input sequence and the target for alignment, BLAST is categorized into the following five primary types:

- **BLASTn**: Aligns nucleotide sequences with nucleotide databases.
- **BLASTp**: Aligns protein sequences with protein databases.
- **BLASTx**: Translates nucleotide sequences into proteins and aligns them with protein databases.
- **tBLASTn**: Aligns protein sequences with translated nucleotide databases.
- **tBLASTx**: Translates nucleotide sequences into proteins and aligns them with translated nucleotide databases.

In this study, to filter duplicate sequences and identify highly similar sequences, we selected the BLASTp tool.

BLASTp is designed explicitly for aligning protein sequences and is primarily used to:

- Identify homologous or functionally related proteins similar to the target protein.
- Predict the functional conservation and potential biological functions of proteins.
- Support protein homology analysis and sequence deduplication.

We integrated the terpene synthase data obtained from UniProt and TeroKit (comprising 9,937 sequences labeled as ‘reviewed’ and 52,282 sequences labeled as ‘unreviewed’) with the aligned sequences from the OMD database (bgc_CDS_ts, a total of 31,259 sequences). This resulted in a final collection of 93,478 sequences, named Collection B.

Subsequently, we will use the makeblastdb command to build its sequence library, Bank A, based on all the sequences updated in BRENDA according to the terpenoid anabolic pathway, Collection

A. Next, the blastp function will be used to compare Collection B with Bank A by running the command as follows:

> blastp -db BankA -query CollectionB -max_target_seqs 5 > blast_results.out

The comparison results are saved in the blast_results.txt file, which details the information about comparing each query sequence against the matching sequences in the database.

The main alignment information is as follows:

First, the Query ID represents the ID of the query sequence, while the Subject ID corresponds to the ID of the matching database sequence. During the alignment process, the similarity between two sequences is measured by the Identity, which is represented as the percentage of identical amino acids between the two sequences. A sequence identity lower than 95% typically indicates significant differences in the two sequences’ nucleotide or amino acid arrangement. This may suggest that the sequences encode different proteins, potentially affecting biological functions. Secondly, low identity could indicate that the sequences originate from other species or evolutionary branches. Additionally, sequence variation may influence the structure and activity of the protein. Lastly, low identity might also reflect genetic mutations adapted to environmental changes. The Alignment length indicates the total length of the participating matches. Mismatches indicate the number of mismatches in the matching, while Gap opens records the number of insertions or deletions during the matching process. In addition, the E-value is used to measure the significance of the matching results, with smaller values indicating more significant and reliable matches, and the Bit score indicates the score of the matching, with higher values meaning better quality of matches. We use the Identity to filter duplicates.

To efficiently filter out sequences with high similarity, we counted the number of lines in the blast_results.txt file (2,500,282) and wrote the Python script remove_95blast.py. The script extracts sequences with a similarity ≥ 95% to the query sequence and stores their IDs in a set. The process involves the following steps: The script begins by reading a new sequence file in FASTA format. It checks whether each sequence’s ID is included in similar sequences. If a sequence’s ID is not found in this set, the sequence is added to a list of unique sequences. Finally, the script outputs the sequences with a similarity below 95%, which are categorized as Collection C. This collection contains 92,076 sequences and is saved in FASTA format.

We combined Collection A with Collection C, resulting in a total sequence library of 456,142 sequences. To ensure the library’s usability for further alignment analysis, we created a final sequence library using the makeblastdb command.

#### 2.1.2 Database construction

##### 2.1.2.1 Structure and function of the database

The structural design of this study in building a terpene synthase database includes three key modules: sequence library, annotated sequence library, and search-match system.

In the data acquisition part, we have completed the acquisition and integration of sequence data from multiple data sources (such as BRENDA, UniProt, TeroKit, OMD, etc.), and all of these data are deduplicated, compared and filtered, and finally merged into a total BLAST sequence library. The sequence library is the core part of this database, which is a wide source of data, storing sequence information related to terpene synthases from different databases, which have been de-emphasized by comparison methods such as BLAST and integrated into a total BLAST sequence library for subsequent matching and querying. Each record in the library contains basic information such as the sequence identifier (e.g., UniProt ID), the sequence itself, the source of the species, etc., which provide a key foundation for subsequent data processing and analysis. Functionally, the sequence library provides basic data support for sequence comparison and functional annotation. As an efficient data query tool, it can help users quickly retrieve and locate the target sequences and further carry out homology analysis, functional prediction, and biological research. The design of this module enables users to find relevant information quickly in the massive data, significantly improving the efficiency of data processing and analysis.

Based on the existing sequences, we mined the information of each sequence in depth. The Annotated Sequence Library stores all the results of functional annotation of sequences by tools such as InterProScan. InterPro is a comprehensive database designed to provide functional and structural domain annotations for protein sequences [13], providing detailed functional descriptions and structural domain information for each protein by integrating information from multiple databases, including Pfam, SMART, and ProDom. InterProScan is a powerful tool for this database, capable of identifying functional regions of proteins through sequence alignment and the Hidden Markov Models (HMM) [14]. By inputting FASTA files, InterProScan generates annotation results in various formats, and the standard output formats are TSV, XML, GFF, and JSON, among which JSON format is widely used in this study. The data in the annotated sequence library provides a detailed functional description for each sequence, revealing information about the sequence’s possible biological functions, related protein families, structural domains, etc. However, it should be noted that the number of sequences per file in the sequence annotation results output by InterProScan is reduced by less than 2000 sequences. This phenomenon primarily arises from several factors. Firstly, the database utilized by InterProScan may lack relevant information corresponding to the input sequences. As a result, some sequences may not successfully identify the corresponding structural domains or functional annotations, leading to unrecorded results. Secondly, InterProScan establishes specific E-value or similarity thresholds during the matching process. If the match between the input sequence and the features in the database falls below these thresholds, the results may be disregarded [15]. In addition, if the FASTA file contains duplicate sequences, InterProScan may report only one match, reducing the number of output results. Finally, some sequences may not have known functional characteristics, so the corresponding annotation results will not be generated. This module provides critical data support for subsequent functional analysis, enabling rapid retrieval of functional information for each sequence and in-depth biological studies of the function of related proteins in the genome. The process is shown in Fig. 2.

**Fig. 1.**
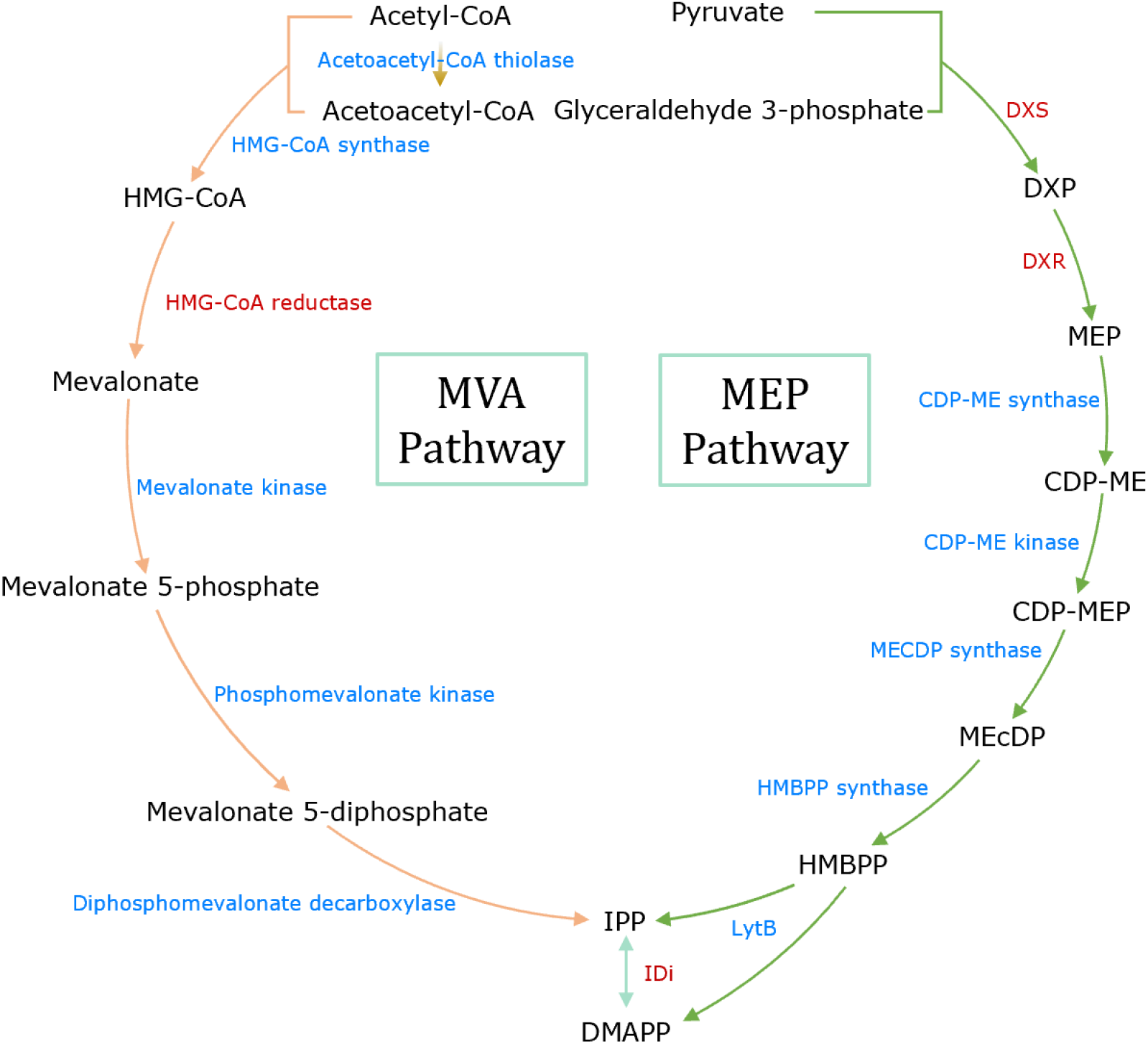
Pathways for the synthesis of terpenoids.

**Fig. 2.**
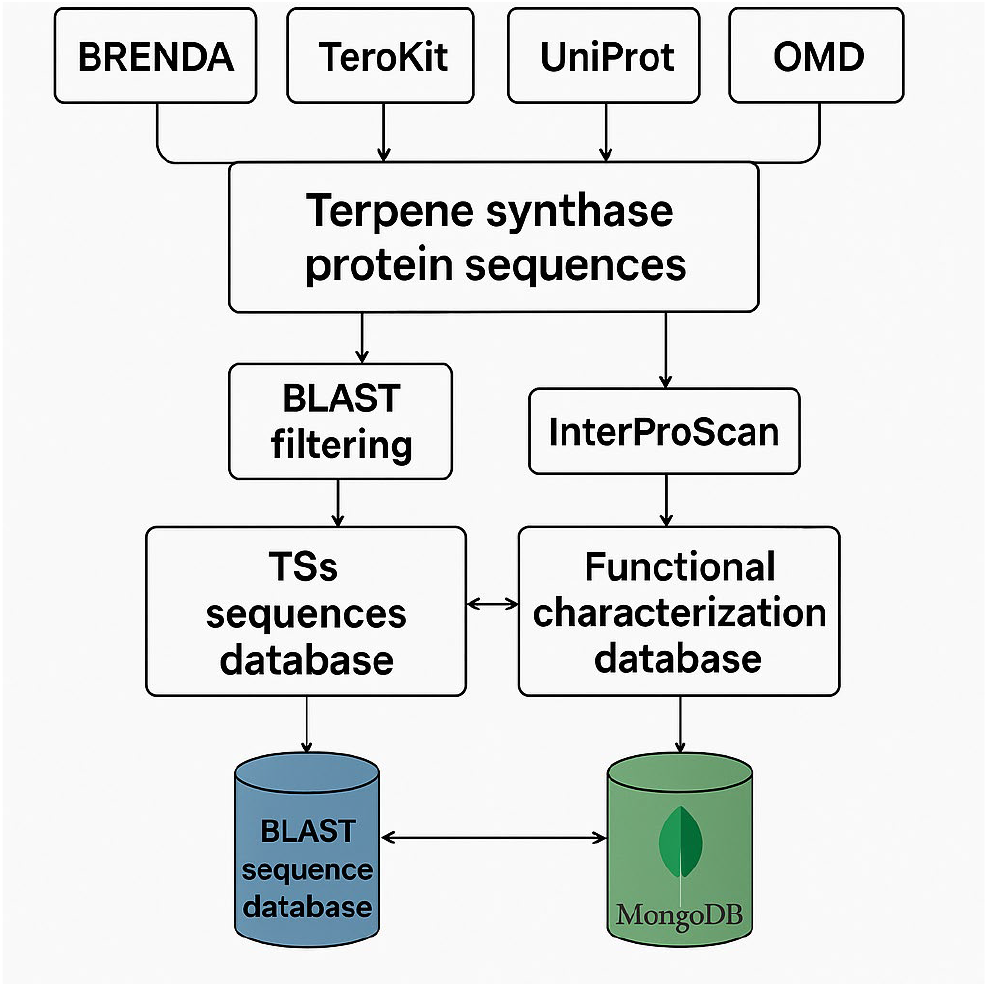
Flowchart of terpene synthase database construction.

To efficiently obtain the annotation information of terpene synthases, we constructed a search system based on BLAST and customized scripts, which are mainly of two types: Shell and Python. First, we wrote a Shell script blastmatch.sh to perform a BLAST comparison of protein sequences for the terpene synthase sequences to be screened and to extract the IDs of similar sequences. During the comparison process, the user can adjust the similarity threshold as needed. When using this script for terpene synthase screening, the user can query the matching status of sequences in a FASTA file with known terpene synthase sequences in the database using the command sh blastmatch.sh xxx.fasta or sh blastmatch.sh “specific sequence.” The output will be in TXT format. The input sequence ID (or name) is displayed on the left side, and the matching ID is shown on the right side. Subsequently, we developed two Python scripts, idmongodb_allanno.py and idmongodb_allandbriefanno.py. Both scripts read the TXT file generated from the BLAST comparison, extract the matched IDs, and query the corresponding annotation information from the MongoDB database. idmongodb_allanno.py outputs the full detailed annotations, while idmongodb_allandbriefanno.py generates both the full detailed annotations and a summary of the annotations in two separate files. Finally, the annotation results are organized and output as CSV format files. To facilitate data processing, each annotated record in the CSV file was categorized according to the source database, and relevant functional information was extracted according to the focus of each database.

During testing, we used the first two sequences from the total sequence library as samples, named ts_first2.fasta, and generated the match result file ts_first2_matchid.txt with the command sh blastmatch.sh ts_first2.fasta. Subsequently, the output file ts_first2_matchid.csv was generated by running the idmongodb.py script to generate the output file ts_first2_matchid_mongoanno.csv. To view the CSV file in a Linux environment, we formatted the output using the command column -t - s, ts_first2_matchid_mongoanno.csv | less -S.

##### 2.1.2.2 Data storage and management methods

In this study, the data storage and management methods mainly rely on two core modules, the BLAST sequence library and MongoDB annotation library. Each module assumes a different role, and together, they ensure the integrity of sequence data, efficient retrieval, and accurate management of annotation results.

The BLAST Sequence Library stores and manages the original protein sequence data and supports subsequent sequence screening and comparison. This structured storage not only ensures the consistency of the data during storage but also provides the necessary data basis for subsequent sequence screening, homology analysis, and functional annotation. With the BLAST tool, we can quickly screen sequences similar to the target sequences and output their matching results as text files. This process improves the operability of the data and provides a convenient query pathway for subsequent sequence analysis and biological studies.

The MongoDB annotation database is responsible for storing and managing protein functional annotation results generated by tools like InterProScan, providing efficient data management and support for subsequent functional analysis and queries. InterProScan performs sequence alignment and utilizes hidden Markov models (HMMs) to identify functional regions within sequences, outputting annotation results that include functional descriptions, protein families, domains, and other relevant information. These annotation data are stored in JSON format and imported into a MongoDB database. With its document-oriented storage model, MongoDB offers a flexible architecture that allows the storage of diverse data types without a predefined schema, making it particularly suitable for handling large-scale and heterogeneous datasets[16]. To facilitate subsequent data retrieval and management, we developed a Python script, json_db.py, to store the InterProScan-generated JSON annotation results in the MongoDB database. The script explicitly extracts the UniProt ID and EC number for each sequence. It stores these key pieces of information as independent fields, enabling fast retrieval and subsequent functional annotation analysis.

## 3 Results and Discussion

### 3.1 Contents of the database

TSDB, as a comprehensive resource for TS sequences and annotations, includes amino acid sequences from several species, TS superfamily and family classifications, protein domains, conserved motifs, binding sites, metabolic pathways, and other related information. TSDB utilizes the BLASTP sequence alignment tool to search for entries in the TSs sequence database that are similar to the target sequence, and based on the alignment results, it annotates the functions of newly identified enzymes. Detailed information about the TSDB is shown in Fig. 3, which illustrates the database’s metadata and functions.

**Fig. 3.**
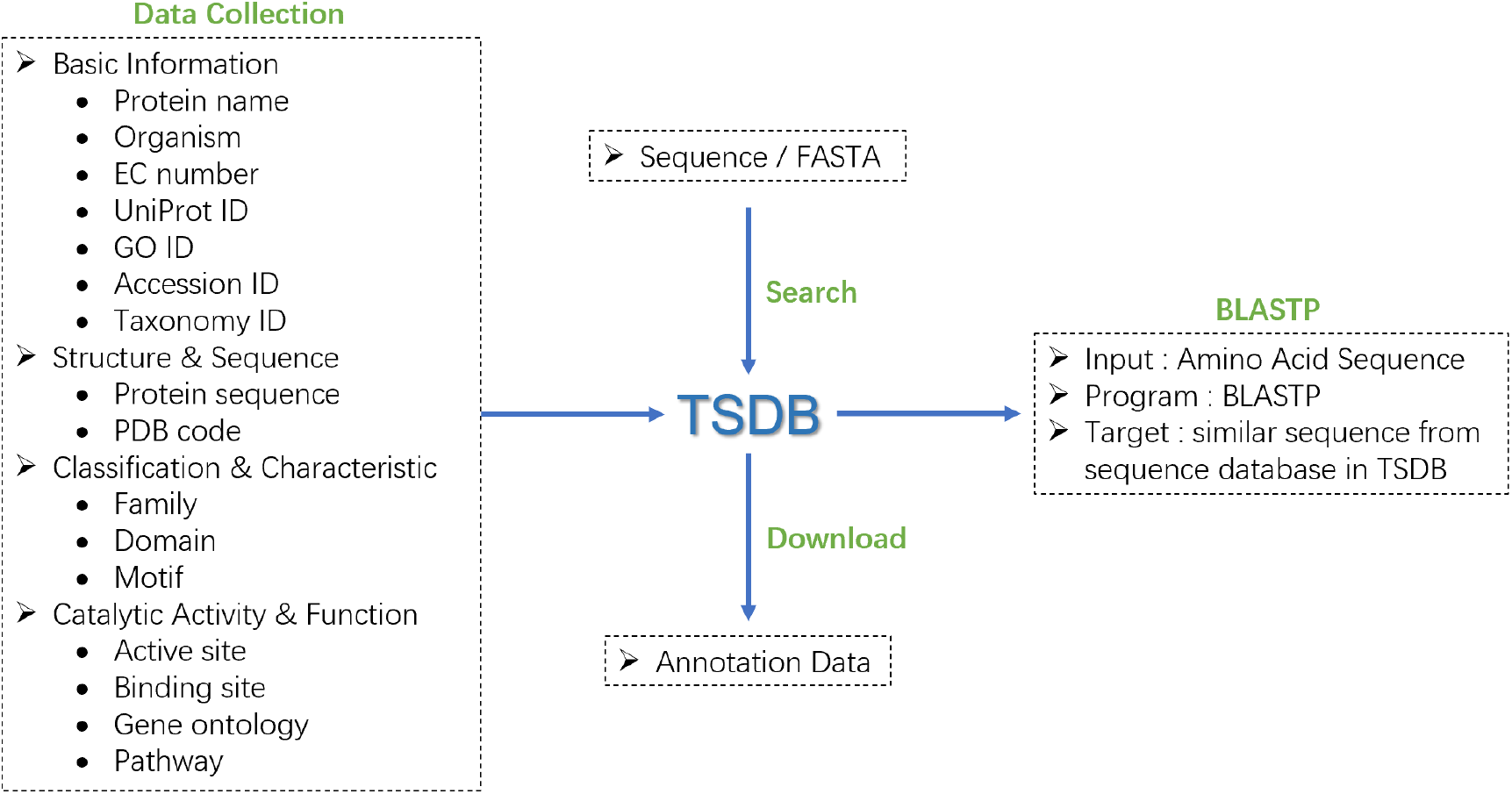
Summary of the TSDB Contents.

Currently, TSDB includes 456,142 amino acid sequences from which corresponding biological source information has been extracted. By comparing with the NCBI Taxonomy database [17], we identified 30,491 unique Taxonomy IDs. Among these, *Bacteria* (25,230) represent the most significant proportion, followed by *Eukaryota* (3,593) and *Archaea* (1,464), while *Viruses* account for only 7 IDs. The detailed species nodes are shown in Table 1.

**Table 1.** Details of species classification nodes in TSDB.

| Superkingdom | Kingdom | Phylum | Class | Order | Family | Genus | Species | Taxonomy ID |
| --- | --- | --- | --- | --- | --- | --- | --- | --- |
| Archaea | 1464 | 1457 | 1319 | 1147 | 1023 | 908 | 614 | 1491 |
| Eukaryota | 3593 | 3593 | 3591 | 3589 | 3581 | 3553 | 3445 | 3593 |
| Bacteria | 25230 | 25181 | 24578 | 23597 | 22834 | 21716 | 7383 | 25400 |
| Viruses | 7 | 7 | 7 | 7 | 5 | 4 | 2 | 7 |
| Total | 30294 | 30238 | 29495 | 28340 | 27443 | 26181 | 11444 | 30491 |

Due to the occurrence of multiple sequences belonging to the same Taxonomy ID, the final count of unique Taxonomy IDs is lower than the total number of original sequences. Additionally, 885 biological sources could not be matched with corresponding IDs in the current NCBI Taxonomy database, likely because the species names used for these sequences are aliases or obsolete names, leading to incomplete searches.

In total, 695 distinct complete EC numbers were identified, covering 392,444 sequences in the database, where *others* includes all EC numbers that account for less than 1% of the total. Furthermore, 129 sequences are associated with incomplete EC numbers, as indicated in their FASTA file descriptions. The distributions are shown in Fig. 4.

**Fig. 4.**
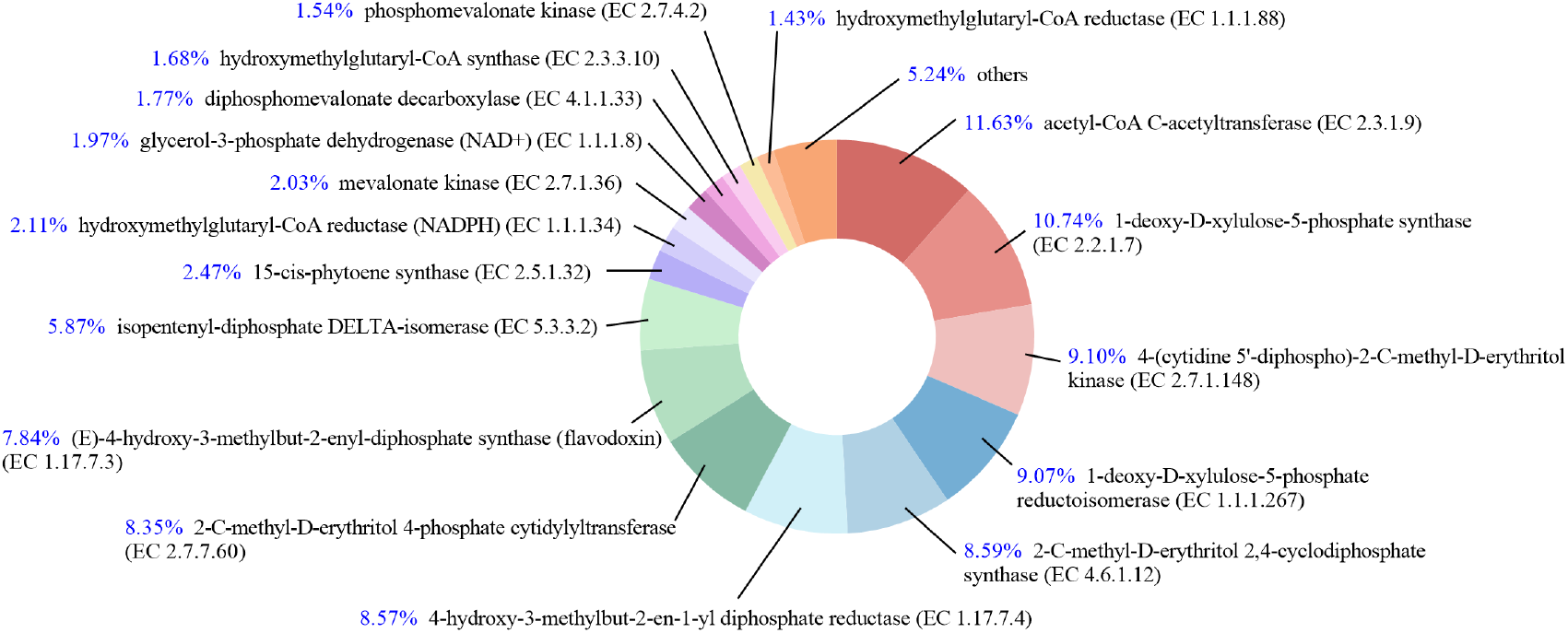
The distributions of terpenoids in TSDB by BRENDA.

### 3.2 Distribution of sequence sources

We conducted a source-based statistical analysis of the 456,142 sequences in the database, and the results indicate that bacterial groups dominate the dataset. The top 20 groups (Fig. 4) reveal that the *Actinomycetia* bacterium and *Gammaproteobacteria bacterium* contribute 2,951 and 2,787 sequences, respectively. In addition, *Helicobacter pylori* and *Chloroflexi bacterium* also represent significant proportions, contributing 2,780 and 2,369 sequences, respectively.

These results suggest that bacterial groups such as *Actinomycetia* and *Gammaproteobacteria* are the most abundant in the database. This may be related to their widespread natural distribution and complex secondary metabolic pathways [18, 19]. This distribution pattern further highlights the database’s representativeness and richness in terms of microbial terpene synthesis potential.

Based on these results, future studies could focus on specific terpene synthase types in these bacterial taxa to reveal their potential roles in natural product synthesis through functional validation and mechanism elucidation. These studies will not only help to reveal the biological basis of microbial terpene synthesis. Still, they will also provide the theoretical basis and data support for expanding the application potential of terpenoids.

**Fig. 4.**
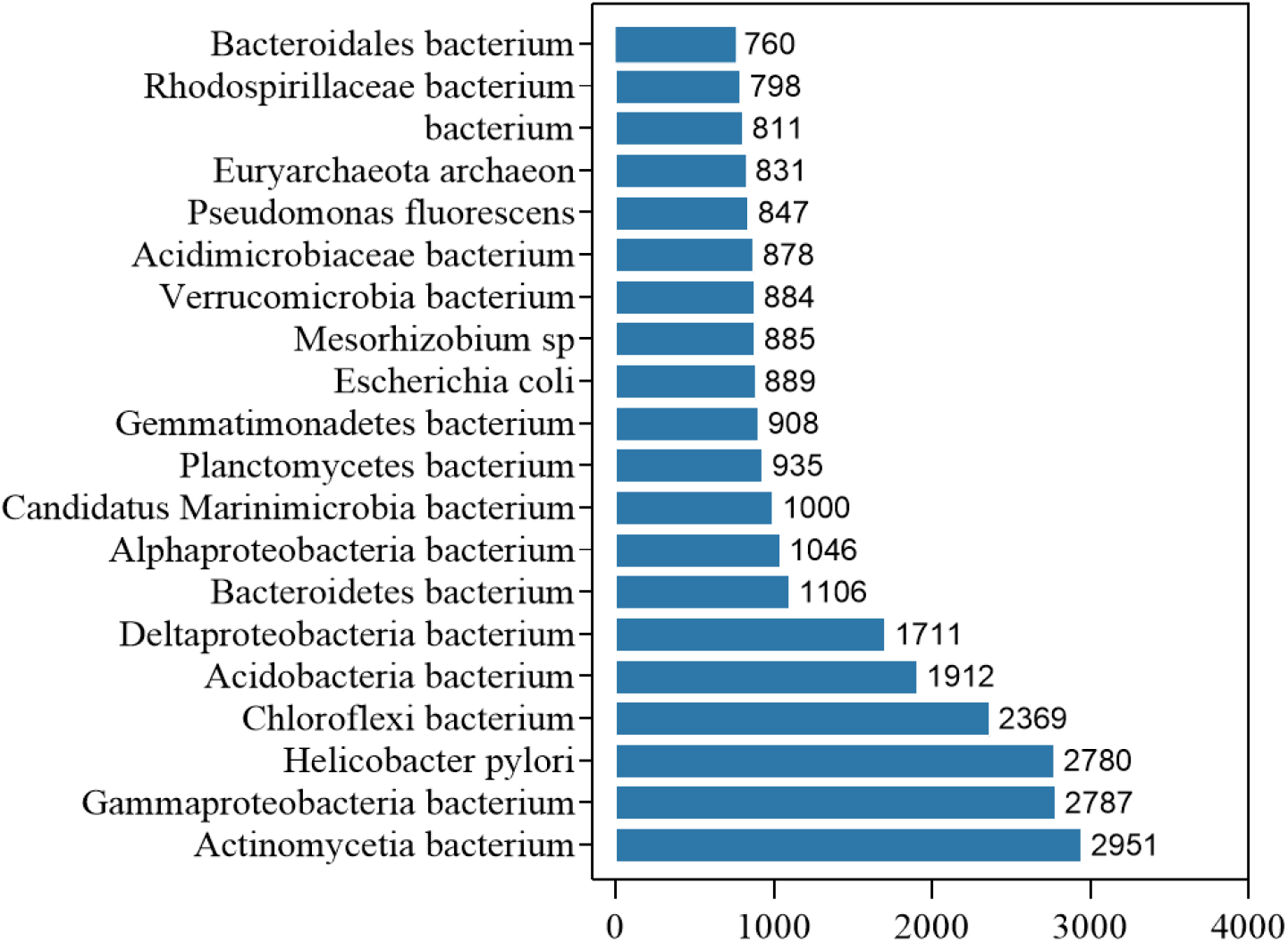
TOP 20 taxa of sequence origin.

### 3.3 Information on functional annotations

TSDB covers 3,499 Gene Ontology (GO) terms, and the distribution statistics of these GO terms reveal functional annotation features related to terpene synthesis. These GO terms span three major categories: Biological Process, Molecular Function, and Cellular Component. The distribution of the top 15 most frequent GO terms is shown in Fig. 5.

**Fig. 5.**
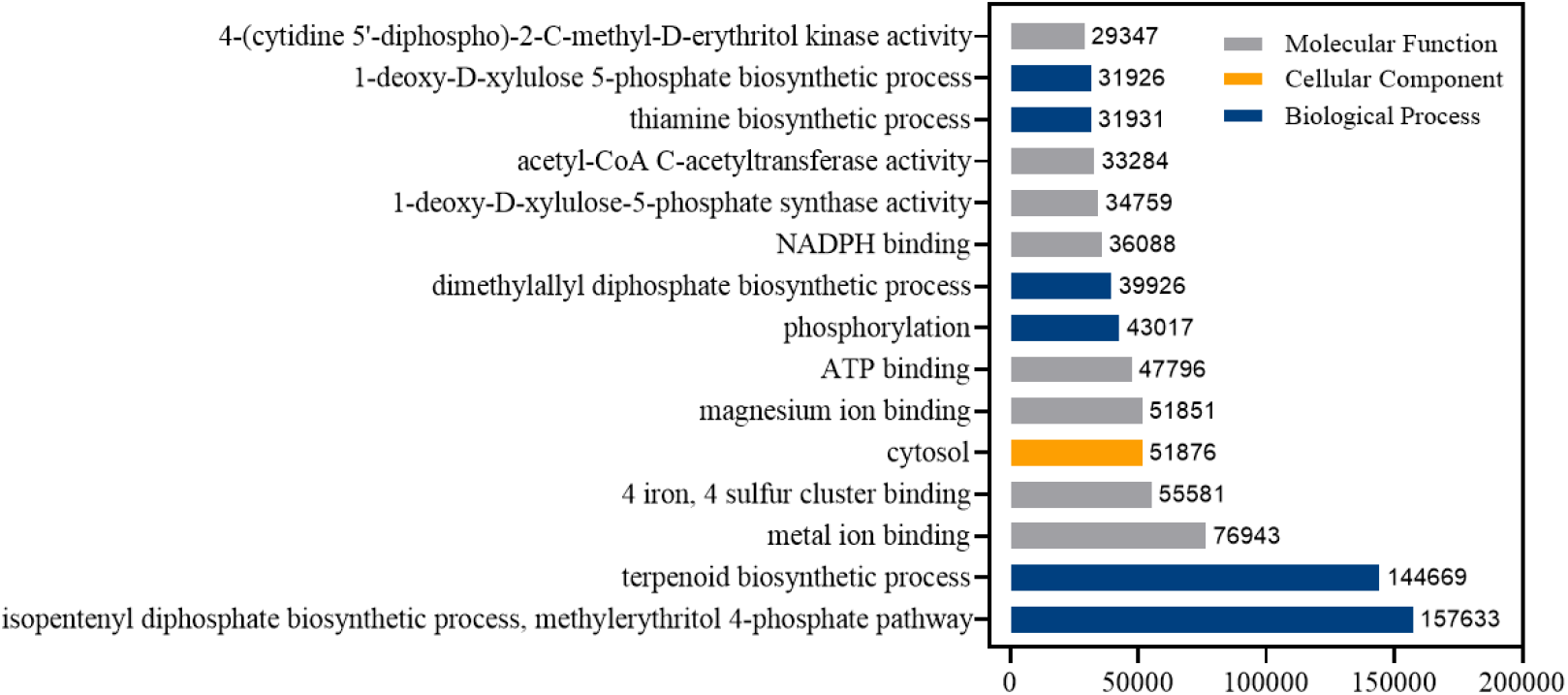
Top 10 GO annotation results in TSDB.

Regarding functional distribution, GO:0019288 (isopentenyl diphosphate biosynthetic process, methylerythritol 4-phosphate pathway) is the most frequently annotated term, covering 157,633 annotation records. This reflects the comprehensive coverage of terpene synthases with isoprenoid metabolism and terpene compound biosynthesis. GO:0016114 (terpenoid biosynthetic process, 144,669 records) and GO:0046872 (metal ion binding, 76,943 records) also represent significant proportions, further revealing the core molecular mechanisms underlying terpene metabolism. Notably, the metal ion binding feature (GO:0046872) supports the critical role of metal ions as cofactors in terpene synthase catalysis. This indicates a close link between enzyme activity and metal dependence and provides valuable insights for future research. Furthermore, the high-frequency annotations of GO:0051539 (4 iron, four sulfur cluster binding, 55,581 records) and GO:0016310 (phosphorylation, 43,017 records) reflect the diversity of terpene synthases in metabolic regulation and electron transfer functions.

### 3.4 Application testing of TSDB

#### 3.4.1 Test methods

To evaluate the annotation effectiveness of TSDB, we used biosynthetic gene cluster (BGC) data from marine microbial genomes as a test set. The testing process begins with FASTA-format genome sequence files, which are then matched against the stored IDs in TSDB using the BLASTP tool. Following this, functional annotation information, including the number of TSs, biological source, terpene production type, and related functional annotations (such as GO terms, protein domains, binding sites, etc.), is extracted based on the matched IDs.

#### 3.4.2 Testing process

In this study, we used gene cluster sequence information downloaded from MIBiG(https://mibig.secondarymetabolites.org/) [20], selecting the test subject with ACCESSION ID BGC0002736.

First, GenBank (gbk) format files containing detailed sequence information for the gene cluster were downloaded from MIBiG. The focus was on CDS (coding sequence) regions, which include terpene synthase sequences involved in the production of terpene compounds. A Python script, gbk2fasta, was used to process the CDS information and extract the amino acid sequences, which were then organized into a FASTA file named BGC0002736_cds.fasta. This test subject contains two CDS regions, with corresponding sequence IDs BGC00027360001 and BGC00027360002.

The FASTA file was used as input for a BLASTP alignment analysis, which identified sequence IDs with over 95% similarity and a bit-score higher than 100 in the TSDB sequence database. The results were saved in a file named BGC0002736_cds_matchid.txt, which lists the input source sequence IDs on the left (BGC00027360001 and BGC00027360002) and the corresponding matching TSDB sequence IDs on the right (P9WEV3 and Q0CRW7, P9WEV2 and Q0CRW6).

Subsequently, using the matching IDs from the BGC0002736_cds_matchid.txt file, the corresponding annotation information was retrieved from TSDB and saved as two CSV files: one detailed file with the suffix mongoanno.csv and one summary file with the suffix mongoanno_summary.csv, to meet users’ different needs.

#### 3.4.3 Annotation results for test objects

To validate the ability of our database to identify terpene-related enzymes, we selected the cluster of known terpene synthesis genes in the MiBiG database with the number BGC0001324 for annotation testing. This gene cluster is derived from the wild tomato species *Solanum pimpinellifolium*, and we obtained its gene cluster structure by antiSMASH (Fig.6).

**Fig. 6.**
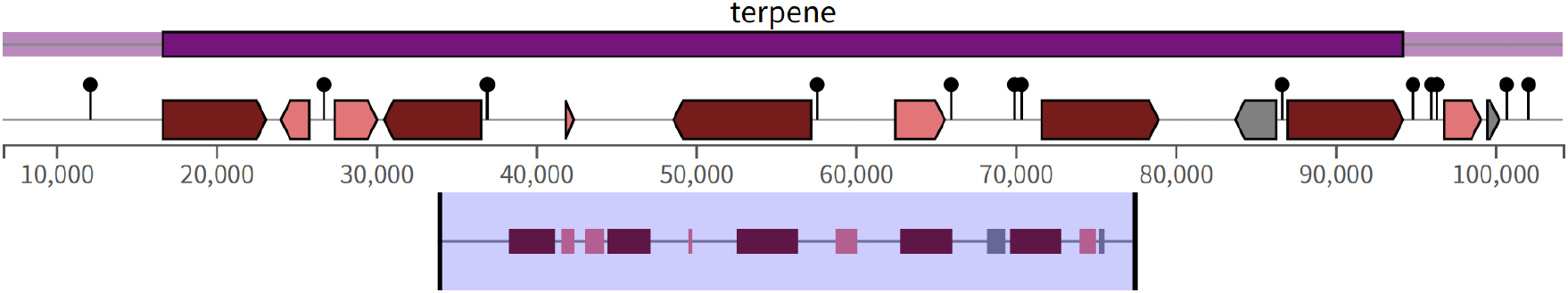
gene cluster structure of BGC0001324.

Through TSDB annotation, we analyzed the well-characterized plant-derived terpene biosynthetic gene cluster BGC0001324 from *Solanum pimpinellifolium*. The core annotation results are illustrated in Fig.7. TSDB identified multiple terpene synthases and associated modification enzymes across various coding sequences (CDSs), consistently assigning them to terpene biosynthetic functions. These results support the presence of a complete and functionally coherent terpene biosynthetic module within the cluster.

**Fig.7.**
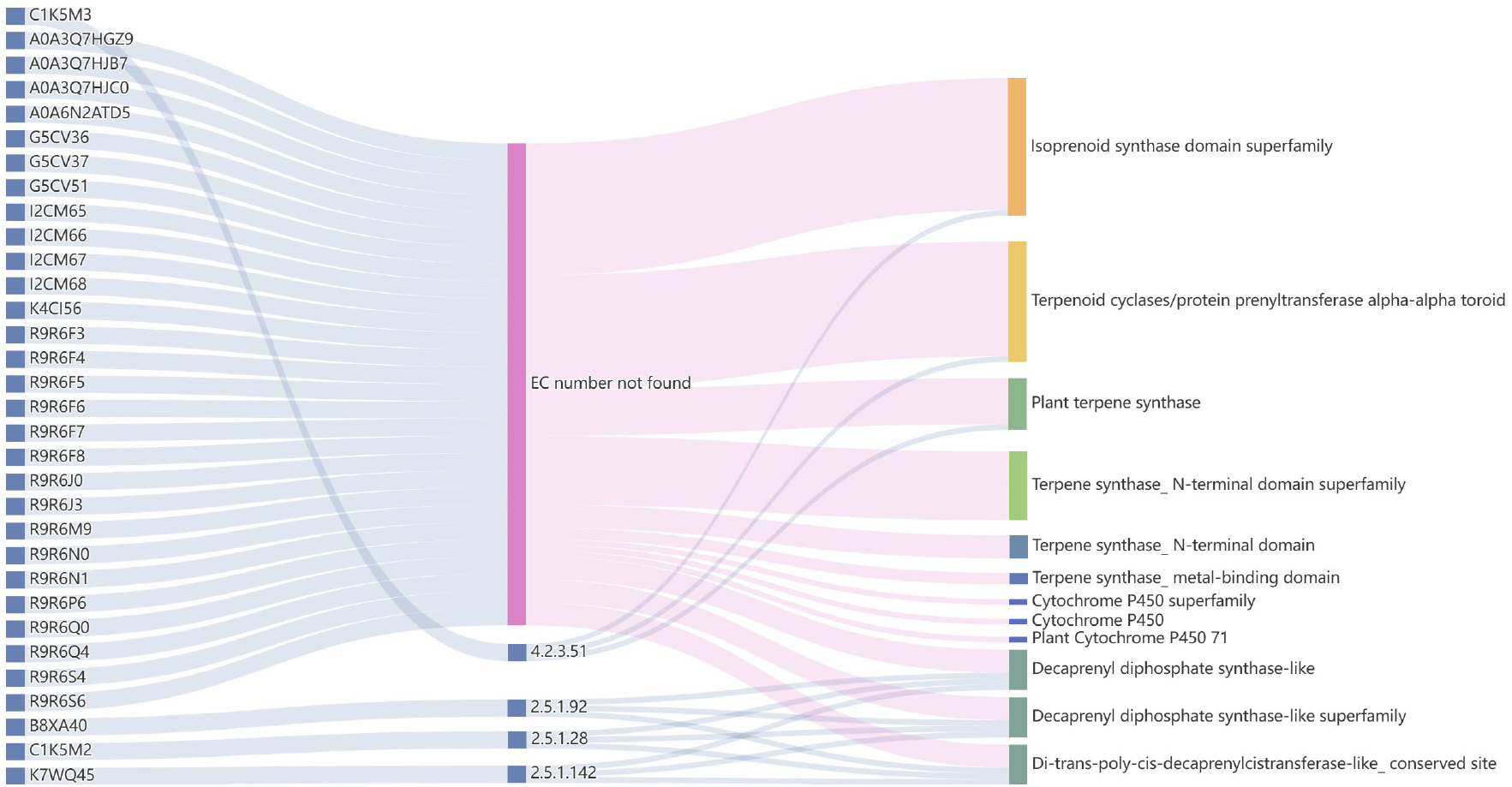
The Sankey of UniProt ID (left), EC number (middle) and signature description (right)

In particular, several CDSs were annotated as terpene cyclases, carrying conserved domain signatures such as Terpene_synth, Terpene_synth_C, and Isoprenoid_synth, all of which are characteristic of class I terpene synthases.[21] These enzymes are involved in catalyzing the cyclization of prenyl diphosphate precursors, such as FPP and GGPP, into diverse terpene skeletons.[22]

In addition to cyclization enzymes, TSDB also annotated several prenyltransferase-like enzymes, including sequences assigned to functions such as Trans-Isoprenyl Diphosphate Synthase (HT), GGPP synthase, FPP synthase, and Isoprenoid synthase type I. These enzymes are known to participate in the elongation of isoprenoid chains, indicating that BGC0001324 contains upstream modules responsible for precursor biosynthesis.[23]

Beyond the core terpene biosynthetic enzymes visualized in the Sankey diagram, TSDB identified auxiliary tailoring enzymes such as cytochrome P450 monooxygenases, which mediate oxidation and hydroxylation reactions that structurally diversify terpene products. These annotations are supported by recurring domain entries such as *P450, CYP503A1-like*, and *Cytochrome P450 family*, further validating their roles in terpene modification.[24]

In addition to enzyme classification, TSDB provides site-level functional annotations (Table 2), offering further structural and mechanistic insights. For example, the protein mapped to UniProt ID K4CI56 was annotated with multiple heme-related sites, including a *heme-binding site*, an *iron (heme axial ligand)*, and a *putative chemical substrate binding pocket*. These features are hallmarks of P450 enzymes, which utilize heme cofactors for electron transfer and oxygen insertion, reinforcing their proposed function in oxidative tailoring. Several other CDSs, such as those corresponding to K7WQ45, R9R6Q0, C1K5M2, R9R6S6, B8XA40, and R9R6F8, were annotated with conserved active sites and dimer interfaces, typically observed in prenyltransferase-like enzymes. These residues are critical for catalysis and oligomerization, suggesting these proteins function as decaprenyl diphosphate synthases or related isoprenoid chain elongation enzymes.

**Table 2.**
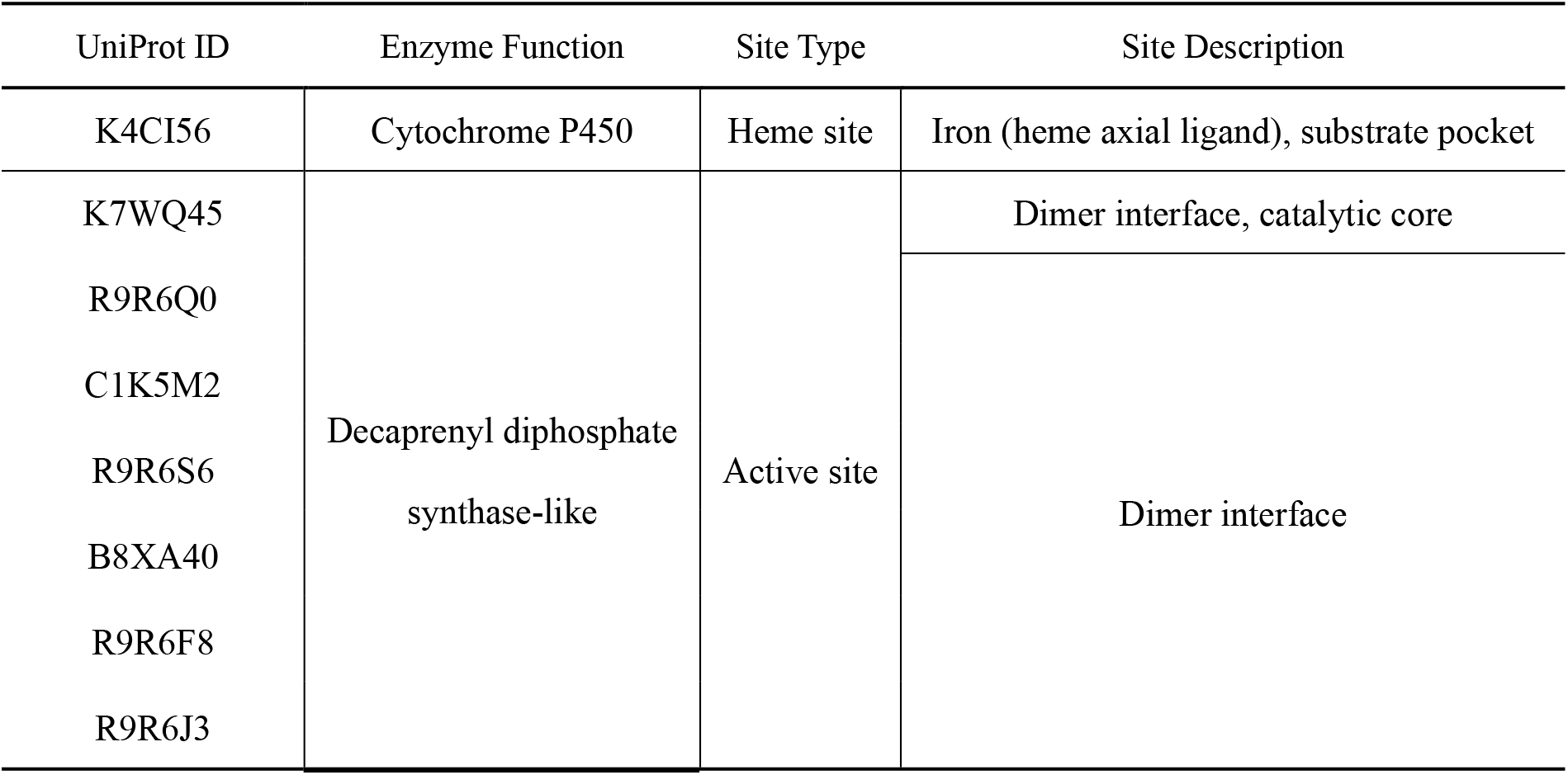
Site-level annotations for proteins encoded within BGC0001324.

These annotations are not only consistent with the functional domains assigned by TSDB, but also reveal the presence of structurally conserved motifs crucial to enzymatic activity. Such detailed annotation enhances the credibility of our predictions and illustrates the capability of TSDB to capture functionally meaningful molecular features beyond standard.

## 4 Conclusions

The development of TSDB marks a significant advancement at the intersection of bioinformatics and natural product research. Unlike existing resources, TSDB systematically integrates functional annotations of MVA/MEP precursor enzymes with terpenoid diversification synthases. By addressing gaps in precursor enzyme coverage and employing domain feature analysis and phylogenetics, TSDB reveals evolutionary patterns in TS catalytic divergence, offering critical insights into the “gene-enzyme-product” nexus. Additionally, its broad taxonomic coverage, particularly in underexplored marine microbes, enhances predictions of terpenoid biosynthetic potential.

Despite progress in annotation depth and species representation, TSDB requires further improvements in dynamic metabolic network modeling and enzyme function validation. Future efforts should integrate experimental data (e.g. enzyme kinetics) to enhance annotation accuracy, develop interactive tools for synthetic biology applications, and employ machine learning to predict enzyme-substrate compatibility. These enhancements will propel TSDB toward systematic and precision-driven terpenoid resource development.

